# Targeted enhancement of flotillin-dependent endocytosis augments cellular uptake and impact of cytotoxic drugs

**DOI:** 10.1101/543355

**Authors:** Farnaz Fekri, John Abousawan, Stephen Bautista, Roya M. Dayam, Costin N. Antonescu, Raffi Karshafian

**Author notes:** Corresponding authors: Costin Antonescu, Department of Chemistry and Biology, Ryerson University, 350 Victoria Street, Toronto, ON, Canada, M5B 2K3, and Raffi Karshafian, Department of Physics, Ryerson University, 350 Victoria Street, Toronto, ON, Canada M5B 2K3.

## Abstract

Cellular uptake is limiting for the efficacy of many cytotoxic drugs used to treat cancer. Identifying endocytic mechanisms that can be modulated with targeted, clinically-relevant interventions is important to enhance the efficacy of various cancer drugs. We identify that flotillin-dependent endocytosis can be targeted and upregulated by ultrasound and microbubble (USMB) treatments to enhance uptake and efficacy of cancer drugs such as cisplatin. USMB involves targeted ultrasound following administration of encapsulated microbubbles, used clinically for enhanced ultrasound image contrast. USMB treatments robustly enhanced internalization of the molecular scaffold protein flotillin, as well as flotillin-dependent fluid-phase internalization, a phenomenon dependent on the protein palmitoyltransferase DHHC5 and the Src-family kinase Fyn. USMB treatment enhanced DNA damage and cell killing elicited by the cytotoxic agent cisplatin in a flotillin-dependent manner. Thus, flotillin-dependent endocytosis can be modulated by clinically-relevant USMB treatments to enhance drug uptake and efficacy, revealing an important new strategy for targeted drug delivery for cancer treatment.

## INTRODUCTION

Conventional drug administration methods such as intravenous injection and oral administration rely largely on diffusion into tumors and cancer cells ^1^. The action of many cytotoxic anti-cancer agents is thus limited by delivery of drug molecules across the plasma membrane of target cancer cells ^2,3^. For example, cisplatin is effective in the treatment of various forms of cancer, and functions by inducing DNA damage and defects in DNA replication once internalized into target cancer cells ^4,5^. However, inefficient cisplatin uptake into cancer cells, toxicity to healthy tissues, and development of resistance limit the effectiveness of cisplatin in the clinic. Advanced drug delivery techniques that can be targeted to cancer cells within tumors can provide a better approach to overcome these limitations and thus improve the efficiency of therapeutic agents such as cisplatin.

The incorporation of drug molecules such as cisplatin into endocytic vesicles contributes to intracellular drug delivery ^6,7^. There are several mechanistically distinct endocytic pathways that operate within cells simultaneously that can be broadly categorized as either clathrin-dependent or clathrin-independent. Clathrin-mediated endocytosis is the principle route of internalization of receptor-bound macromolecules^8^. In contrast, clathrin-independent endocytosis (CIE) encompasses a number of distinct pathways that are diverse with respect to molecular machinery for cargo selection, endocytic vesicle formation and destination of internalized vesicles ^9–11^. Several of these CIE pathways are high capacity and thus can mediate significant uptake of fluid-phase material ^9^. Fluid-phase Internalization is an attractive portal of entry for cancer drugs, as this can ensure the uptake of drug molecules without limitations imposed for specific molecular properties of these drugs (e.g. requirement to bind to specific cell-surface receptors). Therefore, the identification of fluid-phase endocytic mechanisms that can be enhanced for drug delivery purposes and identifying therapeutically-compatible strategies to enhance such an endocytic pathway could provide avenues to achieving more efficient localized drug delivery to cancer cells.

An attractive such CIE mechanism is that delineated by flotillin proteins. The flotillin family is composed of two highly homologous members: flotillin-1 (flot-1 or reggie-2) and flotillin-2 (flot-2 or reggie-1) ^12^. Both members of the family are ubiquitously expressed and highly conserved ^13,14^. Flotillins exhibit cholesterol binding, hydrophobic hairpin insertion into lipid bilayers and acylation, and undergo homo- and hetero-oligomerization to form microdomains enriched in cholesterol and other specific lipids ^15–17^. Flotillin microdomains can serve as scaffolding structures for signaling for a variety of cellular processes ^18^ or to mediate a specific form of CIE ^9,10,19^. Indeed this flotillin-dependent endocytosis can contribute substantially to fluid-phase endocytosis as well as the internalization of cargos such glycosylphosphatidylinositol (GPI)-linked proteins, cholera toxin B subunit, proteoglycans and their ligands and Niemann-Pick C1-like1 (NPC1L1) ^10,20–24^. Flotillin-dependent endocytosis can be modulated by certain cues such as EGF stimulation ^24^.

Identifying cues and their signaling processes that can broadly enhance fluid-phase endocytosis, such as by enhancing flotillin-dependent endocytosis, would be very valuable from a cancer drug delivery perspective. Massive endocytosis (MEND) ^25–27^ may be a particularly attractive mechanism for targeted drug delivery. MEND occurs in response to large intracellular Ca^2+^ transients (e.g. as occurs during plasma membrane perforations) and leads to large increases in fluid-phase endocytosis. While the molecular mechanisms underlying MEND remain incomplete, in fibroblasts MEND requires the enhanced activity of DHHC5 ^26^, a member of family of aspartate–histidine–histidine–cysteine (DHHC) palmitoyltransferases ^28–30^. Upon initiation of MEND, DHHC5 is thought to elicit the broad palmitoylation of cell surface proteins, which then triggers enhanced endocytosis through a poorly understood mechanism.

DHCC5 can palmitoylate flotillin-2 ^31^ as well as neuronal proteins such as postsynaptic density-95 (PSD-95), SynDIG1, GRIP1 and δ-catenin, thus regulating membrane traffic of AMPA-type glutamate receptors in neurons ^32–35^. DHHC5 activity and function is controlled by phosphorylation of DHHC5 on Y533, in a manner dependent on the Src-family kinase Fyn ^35^, which in turn controls DHHC5 cell surface availability and function. Hence, dynamic control of palmitoylation by acyltransferases such as DHHC5 is an emerging signaling system that may be important to control fluid-phase endocytosis, either by direct control of endocytic proteins such as flotillins or by broad palmitoylation of many cell surface proteins (as thought to occur in MEND) to trigger fluid-phase endocytosis.

Leveraging the potential regulation of fluid-phase mechanisms such as MEND-like processes and/or flotillin-dependent endocytosis for drug delivery requires identification of treatments that can therapeutically enhance the action of these endocytic processes. Ultrasound in combination with microbubbles (USMB) ^36–38^ is an emerging strategy for targeted intracellular delivery of drug molecules ^39–42^. Microbubbles (MBs) consist of a gas core with a diameter of less than 5 μm that are encapsulated by lipids, albumin, or polymers, and are systemically injected into the vasculature for clinical ultrasound contrast enhancement ^43^. In addition to this imaging phenomenon, the specific acoustic behaviour of microbubbles in the presence of ultrasound results in a multitude of highly localized effects on nearby cells and tissues ^44^. Microbubbles in circulation are largely inert, and ultrasound can be focused to small (∼ few mm^3^) volumes, making USMB a very attractive strategy to achieve targeted effects in tissues.

USMB treatment elicits the formation of transient pores in the plasma membrane ^44^. An emerging additional outcome of USMB treatment is an increase in the rate of endocytosis ^44–47^. Indeed, we previously reported that USMB enhances the rate of both clathrin-mediated endocytosis and fluid-phase uptake into cells by distinct signaling mechanisms ^48^. However, the molecular mechanisms and intracellular signals by which USMB triggers enhanced fluid-phase endocytosis and whether this phenomenon can contribute to targeted drug delivery to control cancer cell survival had not been addressed, and must be understood in order for this drug delivery strategy to move closer to clinical use. Here, we examine the role of flotillin in the enhanced fluid-phase endocytosis following USMB treatment. We uncovered that USMB treatment elicits a novel signaling pathway that utilizes Fyn and DHHC5 to boost flotillin-dependent endocytosis, thus enhancing fluid-phase endocytosis in several cell types. Importantly, having established this novel mechanism to enhance fluid-phase endocytosis, we uncovered that the enhancement of flotillin-dependent endocytosis by USMB leads to enhanced killing of MDA-MB-231 triple-negative breast cancer cells by cisplatin, highlighting the potential clinical relevance of USMB in targeted drug delivery for cancer treatment by control of flotillin endocytosis.

## RESULTS

### Ultrasound microbubble treatment triggers flotillin-dependent fluid-phase endocytosis

We previously reported that the acid sphingomyelinase (ASMase) inhibitor desipramine synergizes with USMB to enhance fluid-phase endocytosis yet we excluded ASMase as a target for desipramine in the control of endocytosis upon USMB ^48^. Since desipramine is a clinically-approved drug, the possibility of a therapeutic combination of desipramine and USMB could be very promising, so we based our study here of the effects of USMB treatment on fluid phase endocytosis on the combined use of USMB and desipramine.

We first investigated the contribution of flotillin to USMB-induced fluid phase uptake. To do so, we tracked the internalization of the fluid phase marker dextran conjugated to Alexa-488 (A488-dextran) for 30 min after USMB treatment in ARPE-19 (henceforth RPE) cells treated with siRNA to silence flotillin-1 and −2 (**Figure S1A**). RPE cells are ideal to resolve the molecular mechanisms of cell surface phenomena such as endocytosis given their flat morphology. Consistent with our previous observations ^48^, RPE cells transfected with non-targeting (control) siRNA exhibited 12.2 ± 0.1 % and 54.0 ± 0.3 % enhancements in fluid-phase endocytosis upon treatment with USMB or USMB+desipramine, respectively, compared to untreated (basal) cells (n = 3, p < 0.05, **Figure 1A-B**). As we observed previously ^48^, desipramine treatment alone had no effect on fluid-phase internalization. Importantly, cells treated with flotillin siRNA exhibited no change in A488-dextran internalization upon treatment with either USMB or USMB + desipramine (**Figure 1A-B**). These results indicate that flotillins are required for USMB-triggered enhancement of fluid-phase internalization.

**Figure 1.**
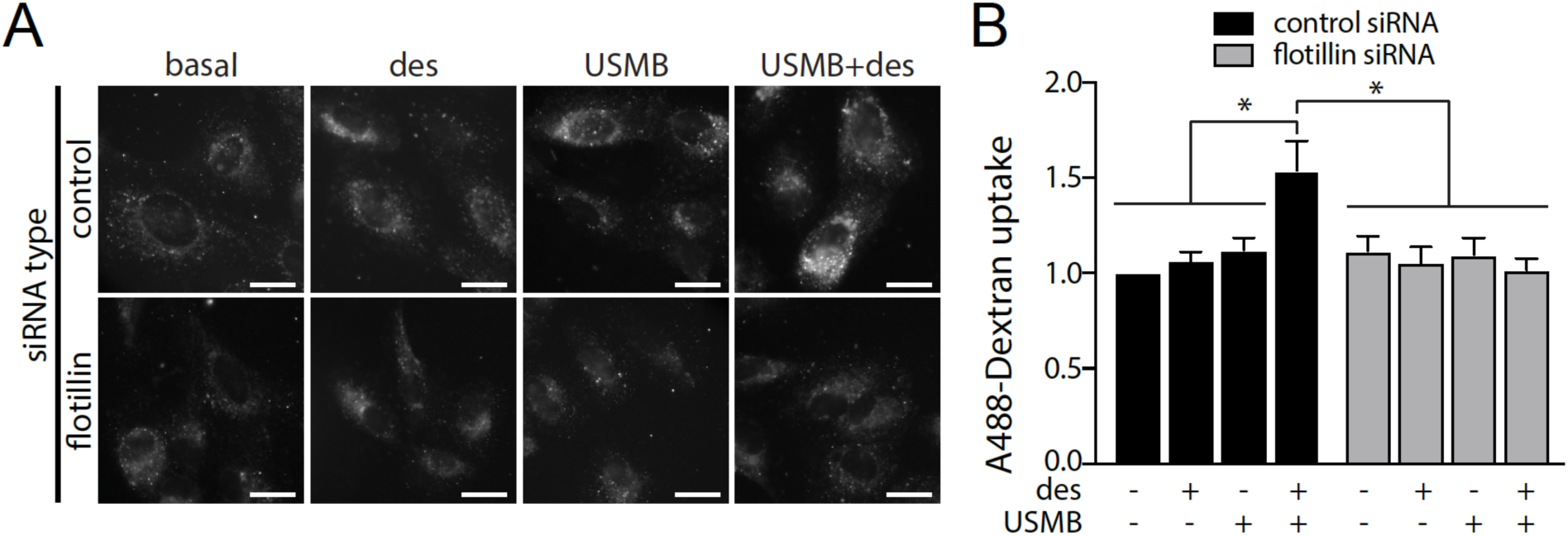
Flotillin is required for fluid-phase internalization triggered by USMB treatment. RPE cells were transfected with siRNA targeting flotillin-1 and −2 (flotillin) or non-targeting siRNA (control). Following transfection, some cells were treated with 50 μM desipramine for 60 min followed by USMB treatment, as indicated, which was subsequently followed incubation with 10 μg/mL A488-dextran for 30 min, fixation, and imaging by widefield epifluorescence microscopy. (A) Shown are representative fluorescence micrographs showing A488-dextran within cells for each treatment, scale 20 μm. (B) Shown are the mean ± SE of total cellular A488-dextran in teach condition n = 3 independent experiments. *, p < 0.05.

The requirement for flotillins in USMB-stimulated enhancement of fluid-phase internalization suggests that USMB may regulate the assembly or dynamics of flotillin structures at the plasma membrane to directly enhance endocytosis mediated by flotillins, or that flotillins may play a permissive role underlying fluid-phase uptake without being regulated by USMB treatment. To distinguish between these possibilities, we examined the impact of USMB on the dynamics of flotillin2-eGFP in RPE cells, monitored by total internal reflection fluorescence (TIRF) microscopy. Of note, flotillin-1 and −2 exhibited nearly complete overlap (**Figure S1B**). As expected ^10^, flotillin2-eGFP formed dynamic fluorescent punctate structures at the cell surface in control cells (**Figure 2A**). We subjected time-lapse image series to automated detection, tracking and analysis of diffraction-limited flotillin2-eGFP structures ^49^. USMB+desipramine treatment did not significantly alter the number (**Figure 2B**) or lifetime (**Figure 2C**) of flotillin2-eGFP structures but decreased the mean fluorescence intensity of flotillin2-eGFP structures (**Figure 2D**). This indicates that USMB+desipramine treatment regulates flotillin structures at the plasma membrane, which may in turn result in enhancement of flotillin-dependent endocytosis.

**Figure 2.**
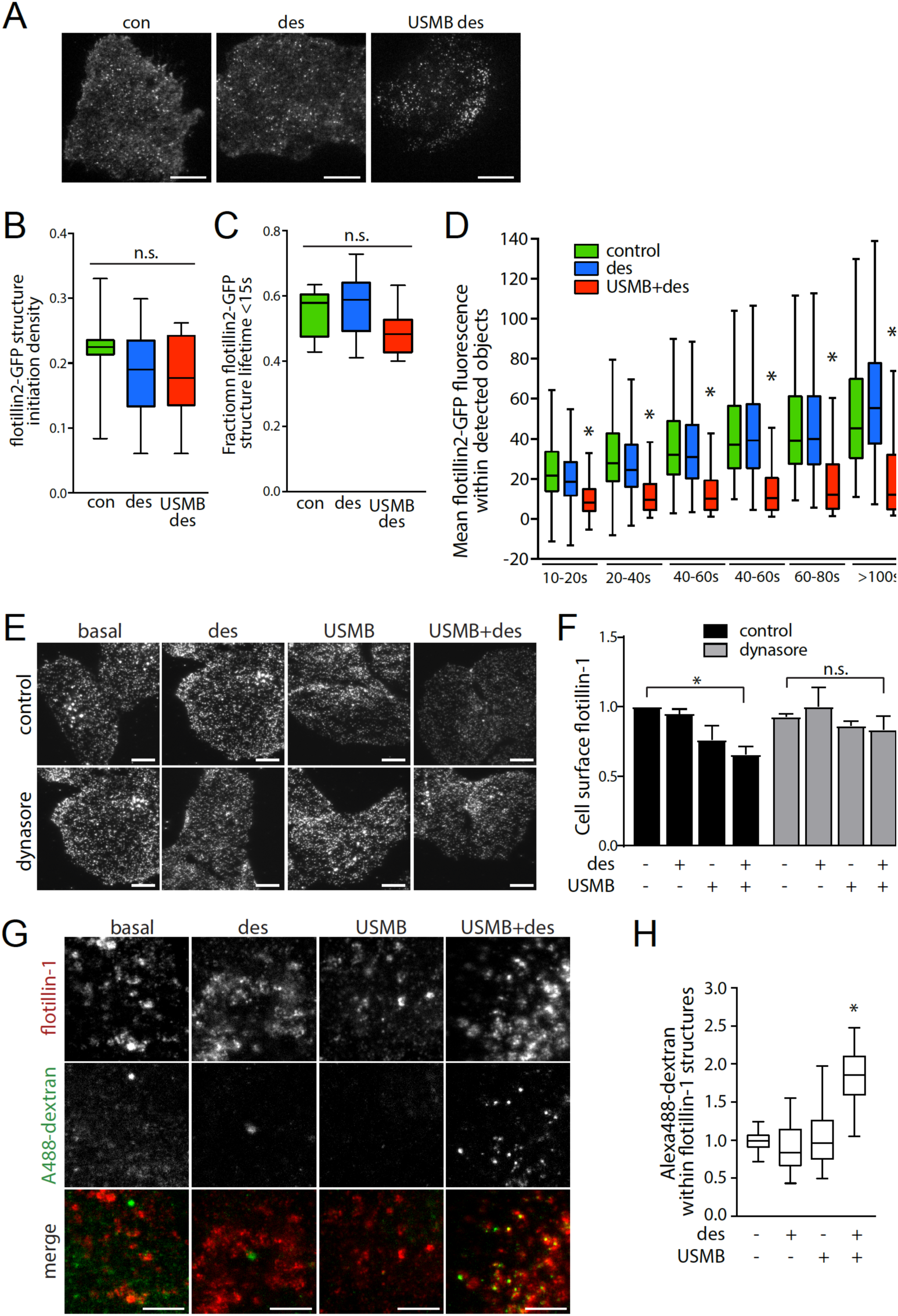
USMB treatment regulates flotillin cell surface levels, dynamics and internalization. (A-D) RPE cells were transfected with cDNA encoding flotillin2-eGFP, then treated with 50 μM desipramine (des) for 60 min followed by USMB treatment, of left untreated (control, con.) as indicated. Cells were then subjected to time-lapse imaging by total internal reflection fluorescence microscopy (TIRF-M), followed by automated detection and analysis of flotillin2-eGFP puncta. Single-frame fluorescence micrographs obtained by TIRF-M are shown in (A). scale 20 μm. The initiation density (B), fraction of structures with lifetimes < 15s and mean intensity (C) of flotillin2-eGFP structures is shown as the median (line) interquartile range (boxes) and full range (whiskers). *, p < 0.05, relative to control. The number of cells (k) and flotillin2-eGFP structures (n) in each condition are as follows: control: k = 8, n = 8626; des: k = 12, n = 9757; USMB+des: k = 11, n = 8272. (E-F) RPE cells were treated with 80μM dynasore and/or 50 μM desipramine for 60 min, followed by USMB treatment or left untreated, as indicated. Cells were then fixed and subjected to immunofluorescence staining of flotillin-1, and then imaging by TIRF-M. (E) Shown are representative TIRF-M fluorescence micrographs, scale 10 μm. (F) Flotillin-1 intensity in each cell in TIRF-M images was quantified to determine cell-surface flotillin-1 levels, and these measurements are shown as mean ± SE. n = 3. *, p < 0.05 (G-H) Cells were treated with 50 μM desipramine (des) for 60 min followed by USMB treatment, of left untreated (control, con.) as indicated. Subsequently, cells were incubated with 10 μg/mL A488-dextran for 30 min, fixed and subjected to staining to detect flotillin-1, and then imaged by spinning disc confocal microscopy. Shown in (G) are representative fluorescence micrographs, scale 5 μm. (H) Flotillin-1 structures were subjected to automated detection and analysis; shown is the mean cellular A488-dextran intensity detected within flotillin structures in each condition, as median (line), interquartile rand (boxes) and full range (whiskers). The number of cells (k) and flotillin2-eGFP structures (n) in each condition are as follows: control: k = 49, n = 26774, des: k = 49, n = 30701, USMB: k = 49, n = 24796; USMB+des: k = 47, n = 28751. *, p < 0.05, relative to control.

Enhanced flotillin-dependent endocytosis will result in decreased abundance of flotillins at the plasma membrane and a concomitant increase of this protein in intracellular compartments^24^. To determine if USMB treatments indeed enhanced flotillin internalization, we next examined the impact of USMB treatment on the plasma membrane localization of endogenous flotillin-1 by immunofluorescence staining and TIRF microscopy, which is highly specific (**Figure S1C**) and allows selective illumination of plasma membrane-proximal fluorophores. USMB or USMB+desipramine treatments reduced flotillin-1 fluorescence intensity in the TIRF field (n = 3, p < 0.05, **Figure 2E-F**). Flotillin-mediated endocytosis requires dynamin for vesicle scission ^50^. Treatment with the dynamin inhibitor dynasore abolished the USMB-elicited reduction in cell surface flotillin-1 (**Figure 2E-F**). These results collectively indicate that USMB treatments trigger enhanced flotillin-dependent endocytosis, consistent with these treatments leading to enhanced flotillin-dependent fluid-phase internalization.

To determine whether the reduction of flotillin-1 from the from cell surface upon USMB treatment indeed contributes to enhanced flotillin-dependent fluid-phase endocytosis, we examined whether USMB stimulation results in overlap of flotillin-1 structures with the fluid-phase marker A488-dextran (**Figure 2G-H**). Discernable localized enrichment of A488-dextran within flotillin-1 structures would only occur upon sequestration of this fluid-phase marker within the lumen of internalized vesicles, which is indeed observed in USMB+desipramine treated cells (**Figure 2G**). To quantify enrichment of A488-dextran within flotillin structures, we subjected these images to automated detection and analysis of diffraction-limited flotillin-1 puncta ^49,51^. USMB+desipramine treatment robustly enhanced the detection of A488-dextran within flotillin structures relative to control samples, or cells only treated with desipramine or USMB alone (**Figure 2H**). Collectively, these observations indicate that USMB treatments enhance fluid-phase endocytosis via regulation that leads to enhancement of flotillin-dependent endocytosis.

### Enhanced flotillin-dependent fluid-phase uptake induced by USMB requires DHHC5

The increase in flotillin-dependent fluid-phase endocytosis by USMB may be related to MEND ^26^, and may thus require the palmitoyltransferase DHHC5. To determine if DHHC5 is required for USMB-stimulated gain in flotillin-dependent fluid phase endocytosis, we next examined the impact of siRNA gene silencing of DHHC5 (**Figure S2A**). While cells subjected to non-targeting (control) siRNA exhibited reduction of 21.5 ± 0.1 % and 45.0 ± 0.1 % in cell surface flotillin-1 levels upon USMB or USMB+desipramine treatment, respectively, cells subjected to siRNA silencing of DHHC5 did not (**Figure 3A-B**). DHHC5 silencing also prevented the gain in fluid-phase endocytosis elicited by USMB+desipramine treatment that is observed in control siRNA treated cells **(Figure 3C-D**). These results indicate that DHHC5 selectively contributes to enhancing flotillin endocytosis and fluid-phase internalization upon USBM treatments.

**Figure 3.**
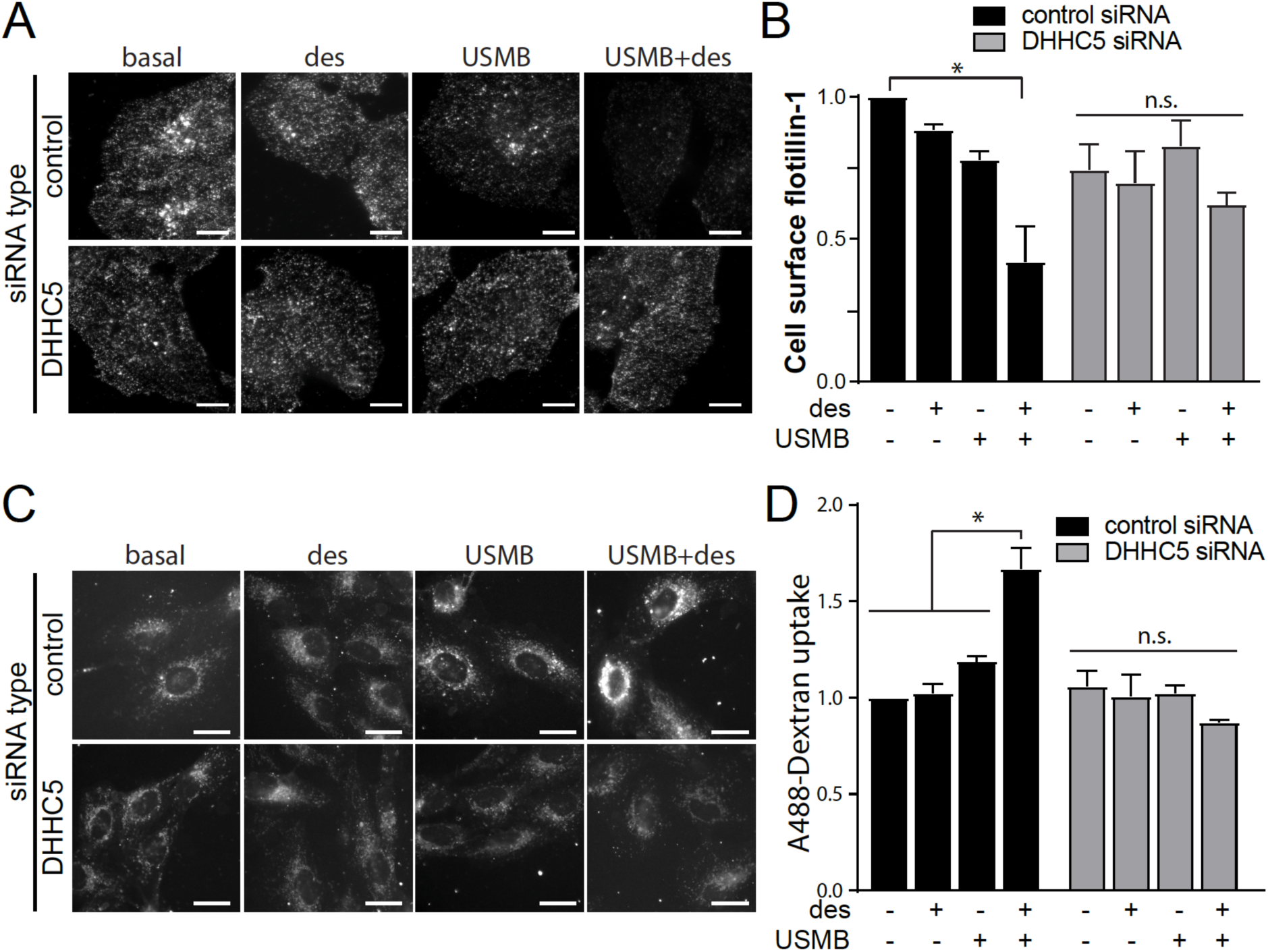
DHHC5 is required for USMB-triggered flotillin and fluid-phase internalization. RPE cells were transfected with siRNA targeting DHHC5 or non-targeting siRNA (control). Following transfection, cells were treated with 50 μM desipramine for 60 min followed by USMB treatment. (A-B) Following treatments and a 30 min incubation, cells were fixed and subjected to immunofluorescence staining of flotillin-1, and then imaged by TIRF-M. Shown in (A) are representative TIRF-M fluorescence micrographs, scale 10 μm. Flotillin-1 intensity in each cell in TIRF-M images was quantified to determine cell-surface flotillin-1 levels, and these measurements are shown in (B) as mean ± SE. n = 3. *, p < 0.05 (C-D) Following treatments, cells were incubated with 10 μg/mL A488-dextran for 30 min, fixed, and imaged by widefield epifluorescence microscopy. Shown in (C) are representative fluorescence micrographs showing A488-dextran within cells for each treatment, scale 20 μm. Shown in (D) are the mean ± SE of total cellular A488-dextran in each condition, n = 3. *, p < 0.05.

To determine how USMB may affect the function of DHHC5, we next examined DHHC5 phosphorylation and cell surface localization. DHHC5 localizes at the postsynaptic membrane of neurons through phosphorylation of Y533 in a manner regulated by Fyn ^35^. To test the possible regulation of DHHC5 by Fyn and/or phosphorylation upon USMB treatment, we first examined the phosphorylation of DHHC5 using phos-tag gel electrophoresis, a method that exaggerates the apparent molecular weight differences elicited by protein phosphorylation ^52^. We observed that USMB and USMB+desipramine elicited a reduction in the apparent molecular weight of DHHC5 observed by phos-tag gel (**Figure 4A**, left lanes), suggesting that USMB induces DHHC5 dephosphorylation. Interestingly, silencing of Fyn (**Figure S3A**) impaired the apparent shift in DHHC5 molecular weight induced by USMB and USMB+desipramine (**Figure 4A**, right lanes). These results indicate that USMB treatment leads to a reduction of DHHC5 phosphorylation, and that this phenomenon is controlled by Fyn.

**Figure 4.**
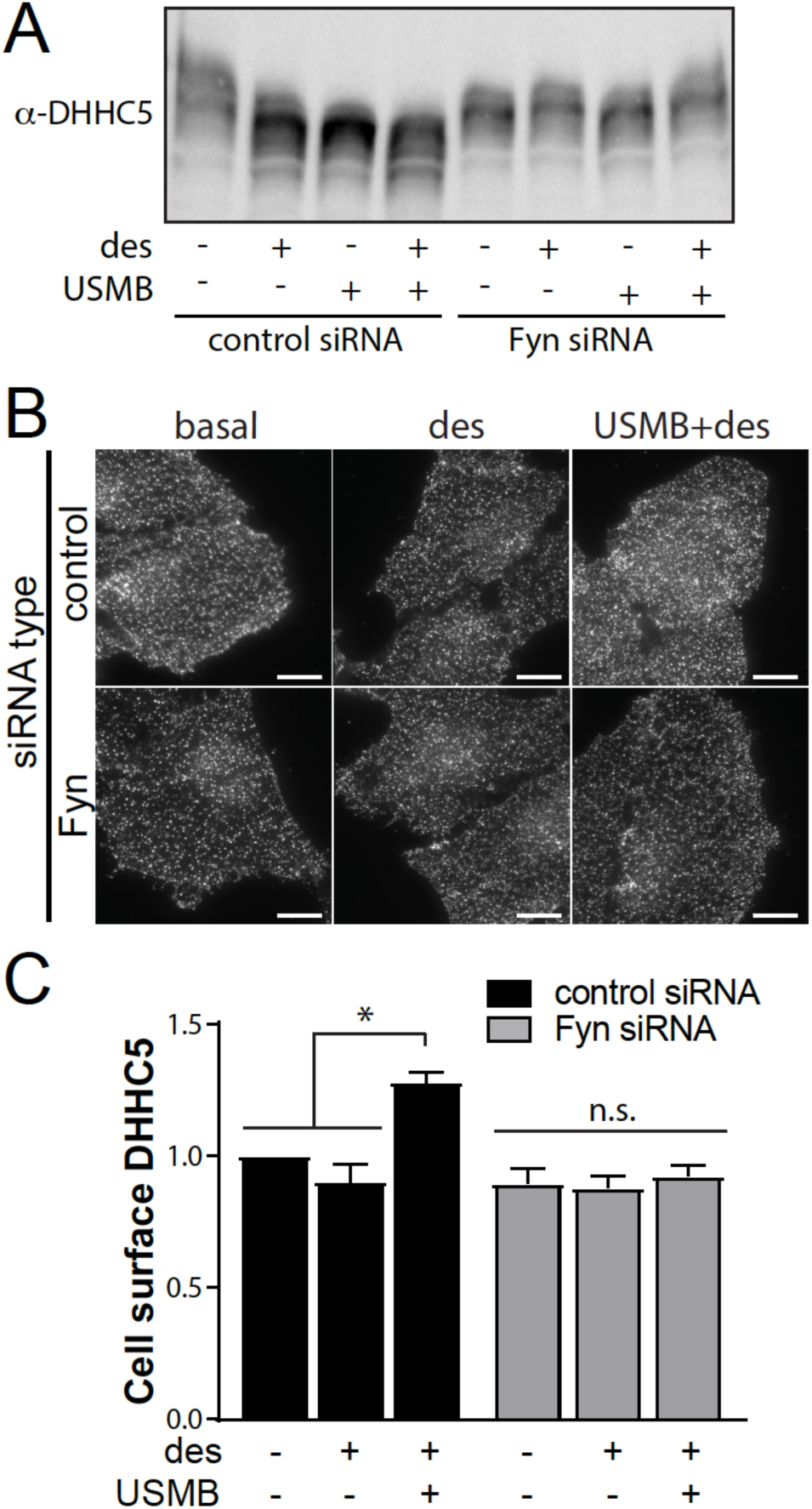
USMB treatment regulates DHHC5. RPE cells were transfected with siRNA targeting Fyn or non-targeting siRNA (control). Following transfection, some cells were treated with 50 μM desipramine for 60 min followed by USMB treatment, as indicated. (A) Following treatments, cells were lysed and whole-cell lysates were resolved by phos-tag SDS-PAGE and western blotting to detect DHHC5. Shown is a representative immunoblot showing DHHC5 staining. (B-C) Following treatments and a subsequent 30 min incubation, cells were then fixed and subjected to immunofluorescence staining of DHHC5, and then imaging by TIRF-M. Shown in (B) are representative TIRF-M fluorescence micrographs, scale 10 μm. DHHC5 intensity in each cell in TIRF-M images was quantified to determine cell surface flotillin-1 levels, and these measurements are shown in (C) as mean ± SE. n = 3. *, p < 0.05.

We next examined how USMB treatment may alter the abundance of endogenous DHHC5 at the plasma membrane using TIRF microscopy. USMB+desipramine treatment increased DHHC5 intensity detected in the TIRF field, and this gain in DHHC5 levels in the TIRF field upon USMB+desipramine treatment was ablated in cells subjected to Fyn silencing (**Figure 4B-C**). This indicates that Fyn controls both the regulation of DHHC5 phosphorylation as well as the gain in cell surface abundance of DHHC5 triggered by USMB treatments.

Phosphorylation of DHHC5 at Y533 controls its cell surface abundance ^35^. We next examined the effect of expression of a phosphorylation deficient Y533A mutant of DHHC5. RPE cells were transfected with wild-type eGFP-DHHC5(WT) or eGFP-DHHC5(Y533A) constructs followed by treatment with USMB, USMB+desipramine or desipramine alone. As expected, cells expressing eGFP-DHHC5(WT) and treated with USMB or USMB+desipramine exhibited decreased flotillin-1 at the cell surface compared to cells not treated with either USMB or desipramine (basal) (**Figure 5A-B,** black bars). In contrast, in cells transfected with eGFP-DHHC5(Y533A), treatment with USMB or USMB+desipramine had no significant effect on the cell surface flotillin-1 levels compared to cells not treated with USMB (**Figure 5A-B,** grey bars). Collectively these results show a functional requirement for regulation of DHHC5 in USMB-stimulated enhancement of flotillin-dependent fluid-phase endocytosis.

**Figure 5.**
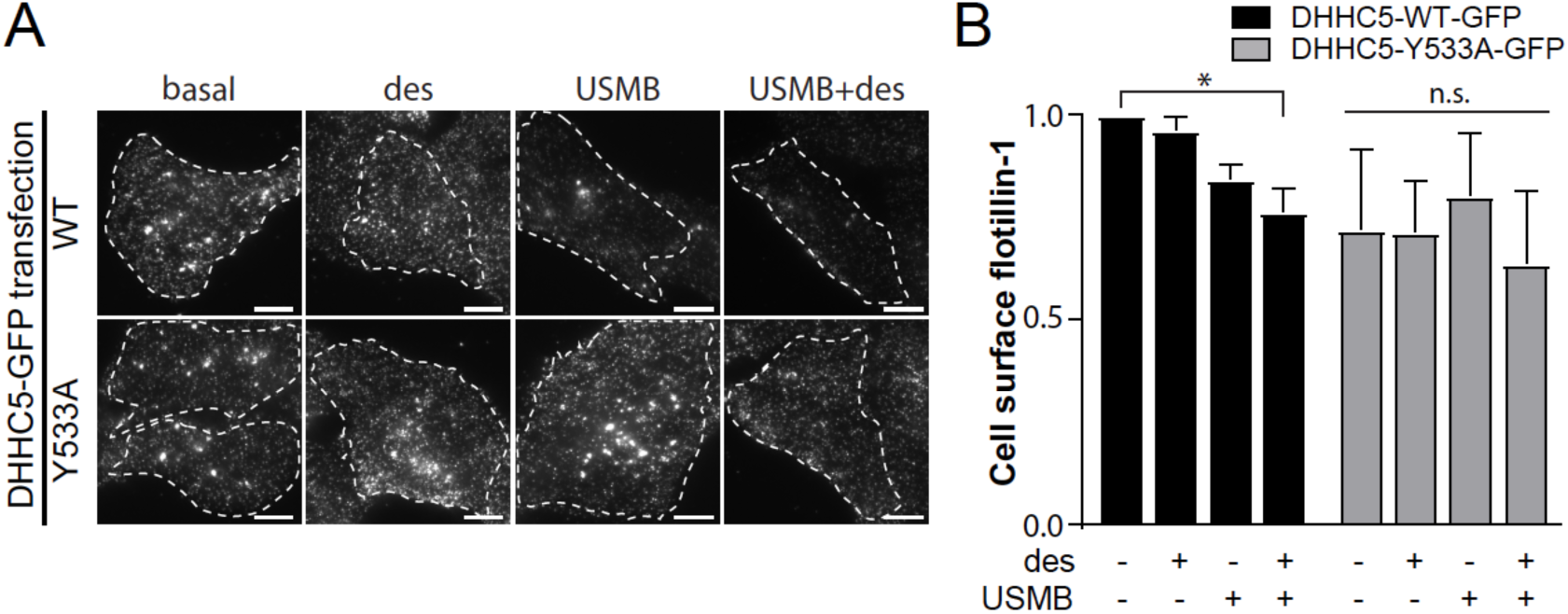
Expression of a phosphorylation-defective mutant of DHHC5 impairs flotillin internalization elicited by USMB treatment. RPE cells were transfected with cDNA encoding either wild-type (WT) or Y533A mutant DHHC5, fused to eGFP. Following transfection, cells were treated with 50 μM desipramine for 60 min followed by USMB treatment. Following a subsequent 30 min incubation, cells were then fixed and subjected to immunofluorescence staining of flotillin-1, and then imaging by TIRF-M. (E) Shown are representative TIRF-M fluorescence micrographs, scale 10 μm. Transfected cells are outlined, and GFP-channel images are shown in **Figure S2B**. (F) Flotillin-1 intensity in each cell in TIRF-M images was quantified to determine cell-surface flotillin-1 levels, and these measurements are shown as mean ± SE. n = 3. *, p < 0.05

### Enhanced flotillin-dependent fluid-phase uptake induced by USMB requires Fyn

We observed that Fyn was required for the regulation of DHHC5 phosphorylation (**Figure 4A**) and for the gain in cell surface DHHC5 (**Figure 4C**) that occur upon USMB treatment. To determine whether USMB-mediated reduction of cell surface flotillin-1 requires Fyn, we used siRNA gene silencing of Fyn (**Figure S3A**). USMB or USMB+desipramine treatment elicited a reduction of flotillin-1 from the cell surface in cells transfected with non-targeting (control) siRNA (**Figure 6A-B,** black bars). In contrast, cells treated with siRNA to silence Fyn did not exhibit a change in flotillin-1 at cell surface upon USMB or USMB+desipramine treatments (**Figure 6A-B,** grey bars), suggesting that Fyn is required for enhanced flotillin endocytosis upon USMB treatment. In addition, cells subjected to siRNA to silence Fyn did not exhibit an increase in fluid-phase endocytosis upon USMB or USMB+desipramine treatments, measured by A488-dextran uptake, as was observed in cells subjected to non-targeting (control) siRNA (**Figure 6C-D**). These results indicate that Fyn is required for the enhanced flotillin-dependent fluid-phase endocytosis elicited by USMB treatment.

**Figure 6.**
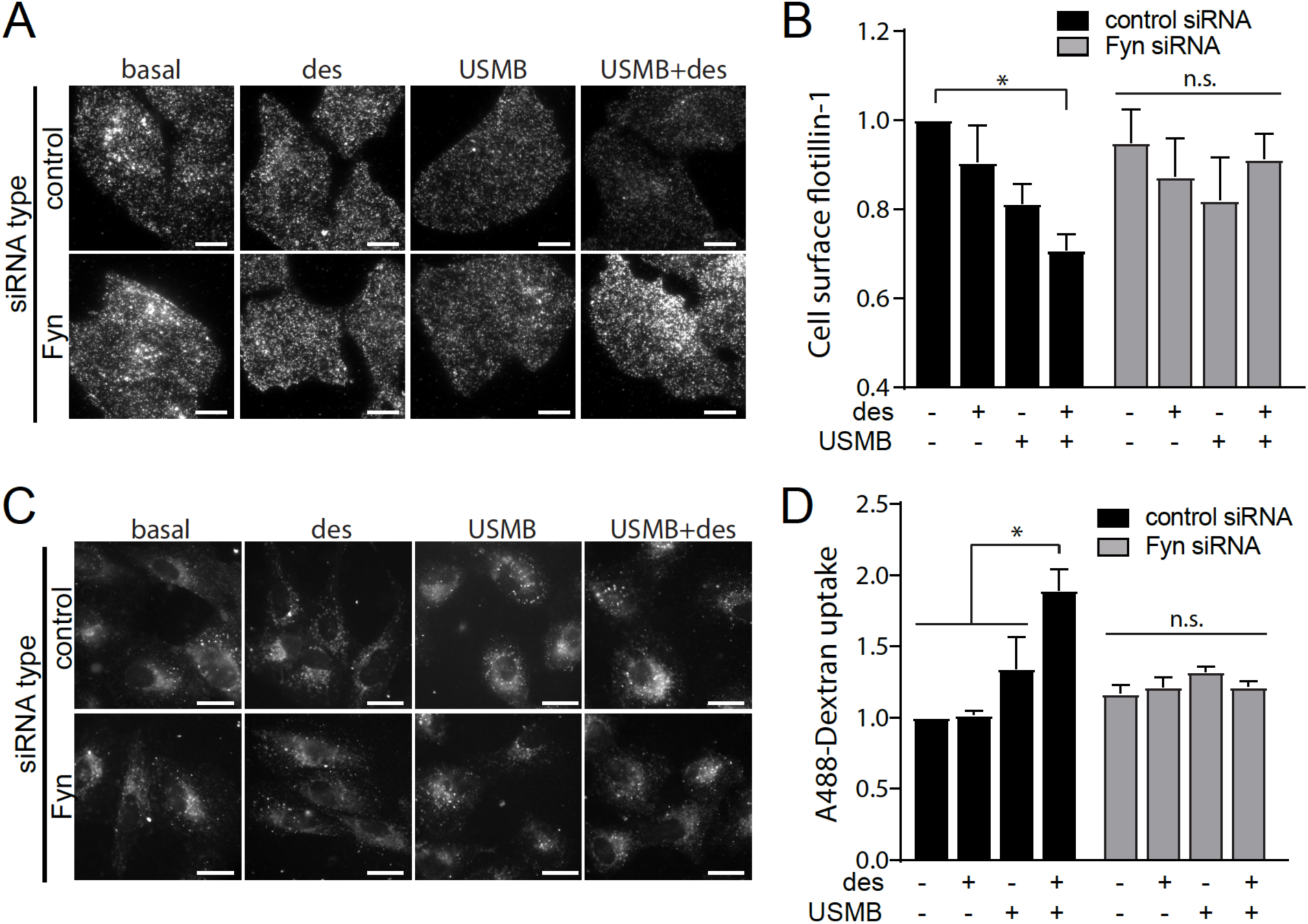
Fyn is required for USMB-triggered flotillin and fluid-phase internalization. RPE cells were transfected with siRNA targeting Fyn or non-targeting siRNA (control). Following transfection, some cells were treated with 50 μM desipramine for 60 min followed by USMB treatment, as indicated. (A-B) Following treatments and a 30 min incubation, cells were fixed and subjected to immunofluorescence staining of flotillin-1, and then imaged by TIRF-M. Shown in (A) are representative TIRF-M fluorescence micrographs, scale 10 μm. Flotillin-1 intensity in each cell in TIRF-M images was quantified to determine cell surface flotillin-1 levels, and these measurements are shown in (B) as mean ± SE. n = 3 independent experiments. *, p < 0.05 (C-D) Following treatments, cells were incubated with 10 μg/mL A488-dextran for 30 min, fixed, and imaged by widefield epifluorescence microscopy. Shown in (C) are representative fluorescence micrographs showing A488-dextran in cells of each treatment, scale 10 μm. Shown in (D) are the mean ± SE of total cellular A488-dextran in teach condition n = 3 independent experiments. *, p < 0.05.

### Enhanced flotillin-dependent uptake induced by USMB enhances cytotoxic drug uptake and action

USMB treatments thus elicit a robust increase in flotillin-dependent fluid-phase endocytosis that also requires DHHC5 and Fyn. To determine how this novel signaling pathway leading to enhanced fluid-phase endocytosis may contribute to enhanced uptake of chemotherapeutic drugs, and thus enhanced ability of these chemotherapeutic drugs to elicit changes in cell viability, we first examined this phenomenon in RPE cells. Following treatment with USMB or USMB+desipramine, cells were incubated in a solution containing 0.03 mM cisplatin (CDDP) for 2 h, followed by washing away of the drug; cell viability was assessed 24 h later (**Figure S4A**). This experimental design allowed for the enhanced fluid-phase endocytosis elicited by USMB treatments to cause an increase in cisplatin uptake immediately (for a 2 h period) following this treatment, and then assessment of the outcome of this initial effect on viability at a later point (24 h later). Importantly, cells treated with USMB+desipramine in combination with cisplatin exhibit a significant reduction of cell viability compared to cells treated with either USMB+desipramine or cisplatin alone (**Figure S4A**).

To determine if USMB+desipramine treatment prior to chemotherapy drug treatment resulted in enhanced uptake and retention of the drug at the time of assessment of cell viability, we repeated similar experimental treatments with USMB and USMB+desipramine, followed by incubation for 2 h in doxorubicin, a cancer chemotherapeutic drug that this intrinsically fluorescent. Cells treated with USMB+desipramine prior to 2 h incubation with doxorubicin exhibited a significantly higher level of cellular doxorubicin assessed 24 h subsequent to drug incubation (**Figure S4B-C**). Together, these results indicate that USMB+desipramine treatment results in enhanced fluid-phase internalization, which contribute to enhanced uptake and action of cytotoxic cancer drugs on cell viability.

To examine how enhanced fluid-phase endocytosis triggered by USMB+desipramine may contribute to enhanced uptake and action of cisplatin in cancer cells, we next examined the impact of this treatment in MDA-MB-231 cells, a model of triple-negative breast cancer ^52^. We used CRISPR/Cas9 genome editing to generate an MDA-MB-231 cell line deficient in flotillin-1 (MDA-MB-231-flot1-KO, **Figure S4D**). While wild-type MDA-MB-231 cells exhibited a robust increase in fluid-phase internalization upon treatment with USMB+desipramine, MDA-MB-231-flot1-KO cells exhibit no discernable change in fluid-phase internalization upon this treatment (**Figure 7A-B**).

**Figure 7.**
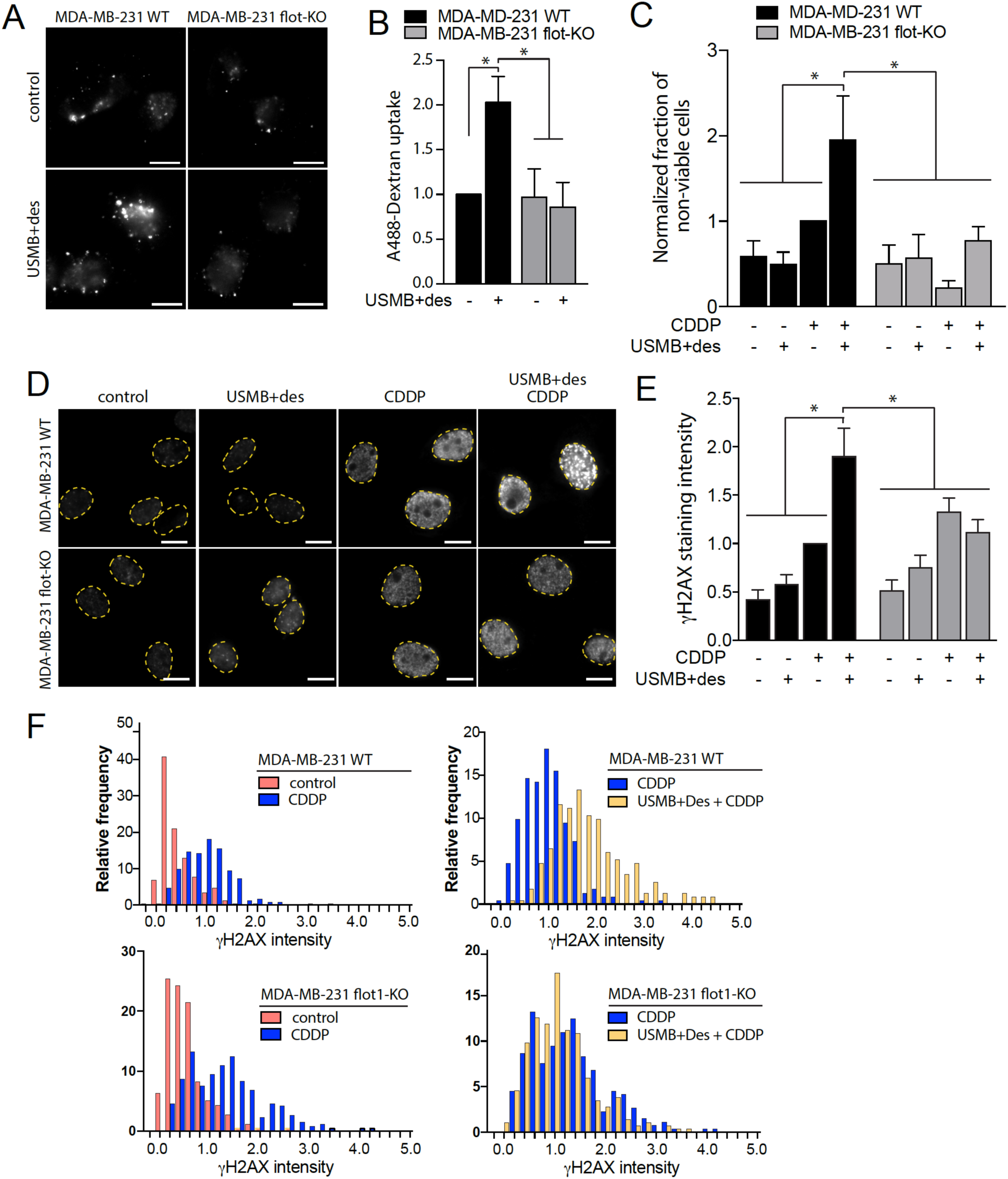
USMB treatments enhance cisplatin-mediated DNA damage and death in breast cancer cells in a flotillin-1 dependent manner. MDA-MB-231 wild-type (WT) or flotillin knockout (flot-KO) cells were treated with 50 μM desipramine for 60 min followed by USMB treatment, as indicated. (A-B) Following treatments, cells were incubated with 10 μg/mL A488-dextran for 30 min, fixed, and imaged by widefield epifluorescence microscopy. Shown in (A) are representative fluorescence micrographs showing A488-dextran in cells of each treatment, scale 20 μm. Shown in (B) are the mean ± SE of total cellular A488-dextran in each condition, n = 3 independent experiments. *, p < 0.05. (C-F) Following USMB and other treatments, cells were incubated with 0.03 mM cisplatin (CDDP) for 2 h (as shown), followed by washing and incubation in growth media (no drugs) (C) 24 h after USMB and/or cisplatin exposure, cell viability was assessed and shown are the mean ± SE of the fraction of non-viable cells, normalized to that of MDA-MB-231 WT cells treated with cisplatin only. n = 3 independent experiments. *, p < 0.05. (D-F) 24 h after USMB and/or cisplatin exposure, cells were fixed and stained to detect γH2AX. Shown in (D) are representative fluorescence micrographs for each condition, scale 10 μm. Shown in (E) is the mean ± SE of γH2AX intensity in each independent experiment. n = 3 independent experiments. *, p < 0.05. Also shown in (F) are frequency distributions of γH2AX intensity measurements in individual cells.

We next examined the impact of this treatment on cisplatin-mediated loss of cell viability. To do so, we again treated cells for 2 h with 0.03 mM cisplatin immediately following the USMB treatment, followed by drug washout and measurement of cell viability 24 h later. Wild-type MDA-MB-231 cells treated with cisplatin only (no USMB+desipramine) exhibited a modest decrease in viability compared to control cells (not treated with either USMB+desipramine or cisplatin) (**Figure 7C**). Importantly, wild-type MDA-MB-231 cells treated with USMB+desipramine prior to cisplatin treatment exhibited a robust increase in cell death compared to either cells treated only with USMB+desipramine or cisplatin alone. Strikingly, MDA-MB-231-flot1-KO cells exhibited no change in viability upon treatment with either cisplatin, USMB+desipramine, or the two treatments combined (**Figure 7C**).

Cisplatin leads to a reduction in cell viability by eliciting DNA damage in the form of double-strand breaks, which first triggers repair mechanisms that can be detected by the presence of γH2AX nuclear puncta ^53^. As expected, in wild-type MDA-MB-231 cells, cisplatin treatment alone elicited an increase in γH2AX levels, which was significantly enhanced by prior treatment with USMB+desipramine (**Figure 7D-E**). Importantly, and consistent with the outcome of cell viability assays, MBD-MB-231-flot-KO cells exhibited no discernable enhancement of γH2AX levels by cisplatin upon prior treatment with USMB+desipramine (**Figure 7D-E**). Collectively, these experiments indicate that the enhanced flotillin-dependent fluid-phase internalization triggered by USMB treatments elicits enhanced uptake, DNA damage and cell death by cisplatin.

## DISCUSSION

We identified that enhancement of flotillin-dependent endocytosis may be a useful strategy for enhancing drug delivery in cancer treatment. This represents one of the first demonstrations that interventions to enhance fluid-phase endocytosis can mediate enhanced drug delivery into cells. The ability to clinically focus ultrasound on small volumes may thus allow targeted drug delivery localized a specific part of a tissue such as a tumor. Moreover, we identified that flotillin-dependent endocytosis triggered by USMB requires the palmitoyltransferase DHHC5 and the Src-family kinase Fyn, and we further identified that USMB induces regulation of DHHC5 that requires Fyn and modulation of DHHC5 phosphorylation. This led us to propose a signaling pathway that links USMB treatment to stimulation of flotillin-dependent endocytosis that initiates with Fyn engagement and control of the palmitoyltransferase DHHC5 (**Figure 8**). Finally, we uncovered that the USMB-stimulated enhancement of flotillin-dependent fluid-phase endocytosis may be significant for enhancing the delivery of drugs such as cisplatin to cancer cells.

**Figure 8.**
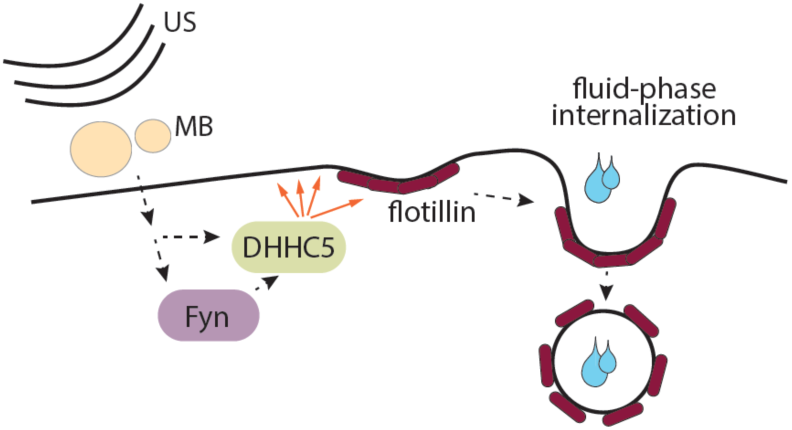
Model of USMB-triggered enhancement of flotillin-dependent fluid-phase internalization. Shown is a diagram depicting ultrasound (US) and microbubble (MB) stimulation, which leads to activation of a signaling pathway involving Fyn and DHHC5, which in turn triggers an increase in flotillin-dependent internalization. This phenomenon elicits an enhancement of fluid-phase internalization, which leads to increased cellular uptake of hydrophilic drugs. In the case of genotoxic drugs such as cisplatin, USMB treatments lead to increased DNA damage and loss of cell viability.

The role of DHHC5 in USMB-triggered enhancement of fluid-phase endocytosis suggests that this endocytic mechanism may be similar to MEND ^26^. MEND can be triggered by a signaling pathway initiated by opening of mitochondrial permeability transition pore channels that leads to DHHC5-dependent enhancement of endocytosis of lipid-ordered membrane domains ^26^, which is consistent with our observation that USMB stimulates flotillin internalization ^48^. Our work is novel in that we find DHHC5-dependent endocytosis to require flotillin, and we demonstrate that stimulation of this pathway can lead to enhanced drug delivery into cells, which had not been examined in previous studies. MEND triggers internalization of ∼50-70% of cell surface area ^26^, which is at least approximately in line with a ∼20-50% enhancement in fluid-phase endocytosis that we observe here.

Our results indicate that USMB treatments may trigger a MEND-like phenomenon, leading to DHHC5-dependent enhancement of flotillin endocytosis. Flotillins themselves are palmitoylated ^15,54,55^, and DHHC palmitoylates flotillin-2 ^31^, suggesting that direct and specific palmitoylation of flotillins by DHHC5 could enhance fluid phase endocytosis upon USMB treatments. This hypothesis is difficult to test experimentally, as perturbation of flotillin palmitoylation by site-directed mutagenesis of flotillins can result in broader alterations in membrane traffic of these proteins ^56^. DHHC5 has several identified substrates and as such it is possible that USMB triggers the palmitoylation of many cell surface proteins, as was suggested to occur in MEND ^26^. Nonetheless, whether USMB elicits broad palmitoylation or palmitoylation of only a few key protein(s) to trigger enhancement of fluid-phase endocytosis, our results strongly indicate that this is flotillin-dependent (**Figure 1**), and results in enhanced formation of flotillin-positive carriers derived from the plasma membrane (**Figure 2**).

How might USMB treatments control flotillin internalization to increase fluid-phase internalization and thus enhance drug delivery? Tracking of flotillin2-eGFP structures in living cells (**Figure 2A-D**) revealed that USMB stimulation led to a reduction of flotillin2-eGFP intensity in these flotillin structures (**Figure 2D**), consistent with USMB stimulation leading to a reduction of flotillin at the cell surface (**Figure 2E**). That USMB treatment decreases the intensity of flotillin2-eGFP in flotillin structures suggests that this treatment may either restrict flotillin oligomerization or may selectively enhance endocytic turnover of larger flotillin structures. This DHHC5-dependent regulation of flotillin-dependent endocytosis adds to the known regulation of flotillin-dependent endocytosis by phosphorylation ^24^. A detailed understanding of the molecular mechanism by which flotillin contributes to endocytosis is currently lacking, which limits more detailed understanding of how USMB treatments and other cues may modulate this process. Future studies thus aimed at elucidating this molecular mechanism will be of great interest.

We identify Fyn and DHHC5 as upstream regulators of flotillin-dependent endocytosis, as perturbations of either DHHC5 (**Figure 3**) or Fyn (**Figure 6**) impair the USMB-stimulated reduction of cell surface flotillin-1 and enhancement of fluid-phase endocytosis (**Figures 4** & **6**). Fyn likely functions upstream of DHHC5 (**Figure 8**) based on the observation that Fyn silencing prevented the USMB-stimulated changes in DHHC5 phosphorylation (**Figure 4A**) and cell surface DHHC5 levels (**Figure 4C**). In neurons, PSD95 recruits Fyn to the cell surface, where Fyn-dependent DHHC5 phosphorylation impairs its endocytosis ^35^. We also observed that Fyn is critical for the regulation of DHHC5 phosphorylation (**Figure 4A**). However, our results suggest that Fyn is required for USMB-induced dephosphorylation of DHHC5 in non-neuronal cells, suggesting additional mechanism by which Fyn controls DHHC5 distinct from direct phosphorylation. Our results are nonetheless consistent with previous reports that Y533 on DHHC5 is critical for the regulation of membrane traffic, likely as a result of phosphorylation of this residue.

Perturbation of flotillin, DHHC5 or Fyn selectively impairs fluid-phase internalization upon USMB treatments, but not in resting cells (**Figures 1, 3 & 6**). Previous studies indeed found that flotillin perturbation impairs fluid-phase endocytosis ^10^, but the contribution of flotillins relative to other forms of clathrin-independent endocytosis varies by cell and condition context ^9^. These results suggest that other endocytic mechanisms are largely responsible for constitutive fluid-phase endocytosis in resting cells. Importantly, flotillin-dependent endocytosis can thus be selectively triggered to boost fluid-phase endocytosis in a selective manner upon USMB stimulation, indicating that this may be an effective strategy for the selective and targeted delivery of drugs into cancer cells.

USMB has been increasingly studied as an attractive and exciting strategy for targeted drug delivery for cancer treatment. At the cellular level, USMB treatment elicits the formation of transient pores, a phenomenon termed sonoporation ^44^. While sonoporation has been proposed as a strategy for enhanced delivery of drugs from the extracellular milieu by simple diffusion, these pores are transient and typically re-seal within < 1 min ^45^. As such, it is these transient pores may only have limited direct contribution to cellular drug uptake. Most experimental strategies that assess the ability of USMB to enhance drug uptake and efficacy involve pre-treatment of cells with the drug in question, and then ongoing incubation in this drug solution following USMB stimulation. Hence, these strategies do not distinguish between access of drugs to the cellular interior through transient pores that form and re-seal during or immediately after USMB (<1 min), or the sustained enhancement of fluid-phase endocytosis as we ^48^ and others ^44–47^ have observed. Here, we added cisplatin starting at 5 min following USMB stimulation, thus allowing us to exclude direct entry through USMB-induced pores as a mechanism for enhanced cellular uptake of cisplatin and enhanced cisplatin-mediated cell killing in USMB-treated cells. That enhanced endocytic uptake via flotillin-dependent endocytosis and not direct entry via USMB-induced transient pores mediates the enhanced cisplatin uptake and action upon USMB treatment is further supported by: (i) the punctate distribution of fluid phase makers upon entry into cells and (ii) that the effect of USMB to enhance cell killing by cisplatin is completely abrogated in flotillin-1 knockout cells (**Figure 7C**).

Our work indicates that USMB, in particular in combination with systemic administration of desipramine, may be an effective combination to selectively enhance drug uptake in cancer cells. This may a particularly effective strategy for targeted drug delivery, as USMB is currently used in the clinic as a diagnostic tool and desipramine is also a clinically-approved agent. Notably, desipramine is without effect on fluid-phase internalization in the absence of USMB, indicating that systemic administration of desipramine and microbubbles, followed by targeted, focused ultrasound can be used for targeted enhanced drug delivery (e.g. to tumors). The identification of the molecular target of desipramine in future studies may extend this proof-of-concept work to additional clinical applications.

The increased flotillin-dependent ability of cisplatin to elicit cell death in cells treated with USMB implies not only that cisplatin is taken up into intracellular vesicles, but also that this treatment results in uptake that has slow recycling of internalized vesicles leading to sustained exposure of cells to drugs within the fluid-phase lumen of these vesicles. Consistent with this, we observed that doxorubicin was retained in USMB-treated cells to a higher extent than in control cells. Moreover, the increased ability of cisplatin to trigger cell death and DNA damage in USMB-treated cells following flotillin-dependent endocytosis suggests that a significant amount of cisplatin must translocate from the lumen of internalized vesicles to access the nucleus (**Figure 7D-E**). The mechanism by which this may occur is unclear and could be due to simple diffusion from the lumen of internalized vesicles retained within the cell or other mechanisms ^4,5^.

In conclusion, flotillin-dependent endocytosis is an endocytic pathway that can be modulated by therapeutically-compatible manipulations such as USMB treatments in order to enhance fluid-phase uptake of cytotoxic drugs, leading to enhanced action of these drugs on cancer cells. This highlights the potential usefulness of USMB or other therapies that control fluid-phase endocytosis in developing more effective cancer treatments, by allowing targeted drug delivery and uptake into tumors or other diseased tissues.

## METHODS

### Materials

Antibodies to detect flotillin-1 and γH2AX were obtained from Cell Signaling Technologies (Danvers, MA) and antibodies to detect flotillin-2 and DHHC5 were obtained from Millipore Sigma (Oakville, ON). Dynasore and desipramine were obtained from Millipore Sigma. Lysine-fixable Alexa-488-conjugated dextran (A488-dextran, 10000 MW) was obtained from Thermo Fisher Scientific (Waltham, MA).

### Cell lines, cell culture and ultrasound treatment

ARPE-19 human retinal pigment epithelial cells (RPE herein) and MDA-MB-231 cells were obtained from American Type Culture Collection (Manassas, VA) and cultured as previously described ^48^. Mycoplasma testing was routinely performed by staining with DAPI, approximately every 2 months. Ultrasound treatment was performed as previously described ^48^. These ultrasound stimulation conditions and activation of and treatment with Definity microbubbles (Lantheus Medical Imaging Inc., Saint-Laurent, QC) were previously optimized and characterized; of note, there was >80% cell viability upon exposure to USMB stimulation ^48,57^.

### Inhibitor and drug treatments

For all experiments, some cells were treated with 50 μM desipramine for 1 h prior to USMB treatment. For the dynamin inhibition experiment (**Figure 2E-F**), cells were either treated with 80 μM dynasore or vehicle control (DMSO) 30 min prior to USMB treatment. For experiments involving treatment with cisplatin (**Figure 7, S4A**) or doxorubicin (**Figure S4C-D**), following treatment with USMB and/or desipramine, cells were incubated for 2 h at 37°C in growth media containing either 0.03 mM cisplatin or 0.03 mM doxorubicin, after which time the solution containing these drugs was removed, the cells extensively washed to remove any non-internalized drug, followed by incubation at 37 C in regular growth media devoid of any drugs or inhibitors.

### Plasmid and siRNA transfections

DNA plasmid transfection were performed as previously described using FugeneHD (Promega, Madison, WI) ^51^. A DNA plasmid encoding eGFP-DHHC5 was a kind gift from Dr. S. Bamji (University of British Columbia, BC) ^35^. This plasmid was used to generate a plasmid encoding eGFP-DHHC5 harbouring a point-mutation corresponding to Y533A by site directed mutagenesis service form BioBasic Inc. (Markham ON). A DNA plasmid encoding flotillin2-eGFP was a kind gift from Dr. G. Fairn (University of Toronto, ON). SiRNA transfections were performed as previously described ^51^, with custom siRNA oligonucleotides (**Table S1**) obtained from Dharmacon (Lafayette, CO).

### CRISPR/Cas9 genome editing

CRISPR/Cas9 genome editing to generate MDA-MB-231 lacking functional flotillin-1 was performed using the Edit-R CRISPR/Cas9 system from Dharmacon (Lafayette, CO), as per manufacturer instructions. Briefly, cells were incubated with a mixture of 0.75 pmoles of Edit-R crRNA targeting flotillin-1 (catalog CM-010636-04-0002), 0.75 pmoles of Edit-R tracrRNA (catalog no U-002005-20), 200 ng of Edit-R hCMV-mKate2-Cas9 DNA plasmid and 15 μL of Dharmafect Duo (Dharmacon) transfection reagent in 300 μL of Opti-MEM media (Thermo Fisher Scientific). Single MDA-MB-231 cells expressing mKate2 were isolated using fluorescence-activated cell sorting; each single cell was grown into separate clonal populations, which were then screened for successful knockout of flotillin-1 by immunoblotting and immunofluorescence microscopy.

### Fluorescent dextran fluid-phase internalization assay

RPE or MDA-MB-231 cells grown on coverslips were treated with desipramine for 1 hour prior to USMB treatments. Following USMB stimulation, cells were treated with 10 μg/mL A488-dextran alone or in combination with desipramine at 37 C for 30 min. After the incubation time, cells were extensively washed and immediately fixed in 4% PFA. Cells were then mounted on glass slides in fluorescence mounting medium (DAKO, Carpinteria, CA).

### Immunofluorescence staining

Endogenous flotillin-1 or −2, DHHC5 or γH2AX were labelled by performing indirect immunofluorescence as previously described ^52^. Samples were fixed in a solution of 4% PFA and permeabilized by Triton-X100 (for flotillin labeling experiments) or ice-cold methanol (for DHHC5 labeling experiments).

### Fluorescence microscopy

Widefield epifluorescence microscopy of fixed samples, as shown in **Figures 1, 3C-D, 6C-D, 7A-B, 7D-F, S1C, and S4B-C** was performed using a 60x (NA 1.35) objective on an Olympus IX83 epifluorescence microscope using a Hamamatsu ORCA FLASH4.0 C11440-22CU camera. Images were acquired using cellSens software (Olympus, Canada, Richmond Hill, ON). Spinning disk confocal and TIRF microscopy of fixed samples was performed using a Quorum (Guelph, ON, Canada) Diskovery combination TIRF and spinning-disc confocal microscope. This instrument is comprised of a Leica DMi8 microscope equipped with a 63×/1.49 NA objective with a 1.8× camera relay (total magnification 108×). Imaging was done using 488, 561 nm laser illumination and 527/30, 630/75 emission filters and acquired using a Zyla 4.2Plus sCMOS camera (Hamamatsu, Bridgewater, NJ). TIRF microscopy imaging of live cells was performed similarly while maintaining cells in temperature (37C) and CO_2_-controlled stage. Time-lapse image series were acquired starting at 5 min after USMB treatment, time lapse images were taken every 60 s for 15 min.

### Image analysis

Quantification of whole-cell intensity of flotillin-1, DHHC5, or internalized fluid-phase A488-dextran, or nuclear γH2AX was performed as previously described ^51,52^ using ImageJ software (National Institutes of Health, Bethesda, MD) ^58^. Detection and analysis of diffraction-limited internalized flotillin-1 puncta (**Figure 1G-H**), was performed as previously using custom software developed in Matlab (Mathworks Corporation, Natick, MA), as described previously for cell surface endocytic structures ^49,59–61^ and internalized vesicles or endosomes ^51^. Detection, tracking and analysis of cell surface flotillin2-eGFP in time-lapse image series obtained by TIRF-M (**Figure 2B-D**) was performed as previously described for cell surface clathrin structures ^49,62^.

### Immunoblotting

Whole cell lysates were prepared in Laemmli sample buffer, resolved by SDS-PAGE and probed with specific antibodies, and the resulting signal intensity quantification were performed as previously described ^52^. To examine phosphorylation of DHHC5 (**Figure 4A**), we used the phos-tag gel system, which results in exaggeration of differences in apparent molecular weight of phosphorylated forms of specific proteins, as previously described ^52^.

### Cell viability measurements

To perform the proliferation/viability assay as shown in **Figure 7B**, 24h following treatments as indicated cells were detached using 0.025% trypsin and cells were counted using a Countess II FL Automated Cell Counter (Thermo Fisher Scientific). The number of non-viable cells was determined by the loss of cells from the initial number of seeded cells and expressed as a normalized value. To perform the cell proliferation/viability assay as shown in **Figure S4A**, RPE cells seeded in 6 well plates were treated as indicated. After 24 h, cells were stained with crystal violet, then solubilized in a solution of 0.1% SDS, followed by measurement of absorbance at 595 nm. Cell viability was expressed as percentage of the crystal violet signal in control (untreated) cells.

### Statistical analyses

Statistical analysis was performed as previously described ^52^. Measurements of samples involving one experimental parameter and more than two conditions (**Figures 2A-B, 2H, S4A, S4C**) were analyzed by one-way ANOVA, followed by Tukey post-test to compare differences between conditions, with p < 0.05 as a threshold for statistically significant difference between conditions. Measurements of samples involving two experimental parameters (**Figures 1B, 2F, 3B, 3D, 4C, 5B, 6B, 6D, 7B, 7C, 7E**) were analyzed by two-way ANOVA, followed by Tukey post-test to compare differences between conditions, with p < 0.05 as a threshold for statistically significant difference between conditions.

## ACKNOWLEDGEMENTS

Funding for this research was provided by a Discovery Grant from the Natural Sciences and Engineering Research Council, a New Investigator Award from the Canadian Institutes of Health Research (CIHR) and an Early Researcher Award (Ontario Ministry of Research, Innovation and Science) to C.N.A., a Discovery Grant from the Natural Sciences and Engineering Research Council to R.K. and an Ontario Graduate Scholarship to F.F and S.B.

## AUTHOR CONTRIBUTIONS

F.F., C.N.A. and R.K. wrote the main manuscript text, F.F., J.A., S.B., and R.M.D. performed the experiments, and F.F., J.A. and C.N.A. prepared the figures. All authors reviewed the manuscript.

## ADDITIONAL INFORMATION

The author(s) declare no competing interests.

