## Supplemental Materials for "Targeted enhancement of flotillin-dependent endocytosis augments cellular uptake and impact of cytotoxic drugs"

This Supplemental Materials file contains

Table S1
Figure S1
Figure S2
Figure S3
Figure S4

| **siRNA name (target)** | **Sense strand sequence** | **Antisense strand sequence** |
|---|---|---|
| non-targeting control | CGU ACU GCU UGC GAU ACG GUU | CGT ACT GCT TGC GAT ACG GUU |
| DHHC5 | CUG UGA AGA UCA UGG AUA AUU | UUA UCC AUG AUC UUC ACA GUU |
| Fyn | AGG AAG AGC UCU GAA AUU AUU | UAA UUU CAG AGC UCU UCC UUU |
| flotillin1 | UGG CCA AGG CAC AGA GAG AUU | UCU CUC UGU GCC UUG GCC AUU |
| flotillin2 | GGA UGA AGC UCA AGG CAG AUU | UCU GCC UUG AGC UUC AUC CUU |

**Table S1. Sequences of custom siRNA oligonucleotides used in this study**. Shown are the sense and antisense sequences for each siRNA oligonucleotide used in this study.

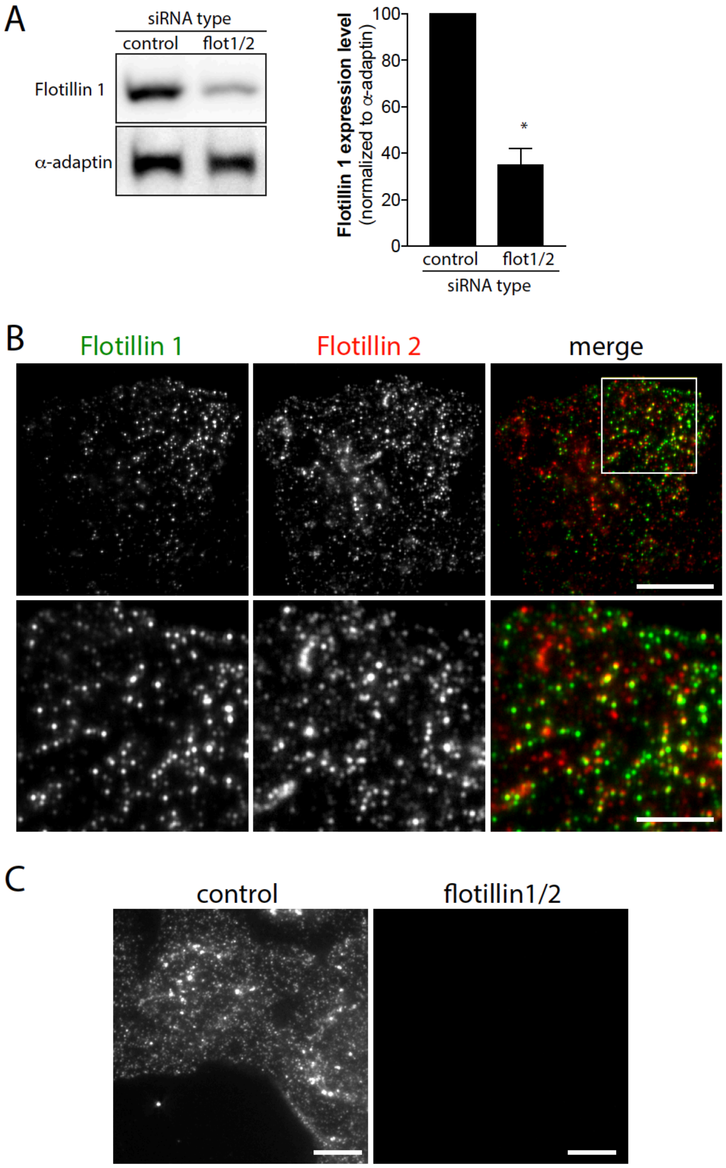


**Figure S1. Flotillin knockdown and immunofluorescence detection.** (A) RPE cells were transfected with siRNA targeting flotillin-1 and -2 (flotillin) or non-targeting siRNA (control). Whole-cell lysates were resolved by SDS-PAGE and subjected to immunoblotting to detect flotillin-1 or alpha-adaptin (AP2, loading control). Shown (left panels) are representative immunoblot and the mean ± SE of flotillin-1 detected by this method (right panels). This demonstrates effective knockdown of flotillin1 by siRNA silencing. (B) RPE cells were subjected to immunofluorescence staining to detect flotillin-1 and flotillin-2 and then subjected to imaging by total internal reflection fluorescence microscopy (TIRF-M). Shown are representative fluorescence micrographs that show the extensive co-localization of flotillin-1 and flotillin-2, scale 10 μm. This indicates that the vast majority of flotillin structures in these cells are positive for both flotillin-1 and flotillin-2. (C) RPE cells were transfected with siRNA targeting flotillin-1 and -2 (flotillin) or non-targeting siRNA (control), followed by immunofluorescence staining to detect flotillin-1, then subjected to imaging by TIRF-M. Shown are representative fluorescence micrographs, scale 10 μm, which indicate the specificity of flotillin-1 staining for flotillin-1.

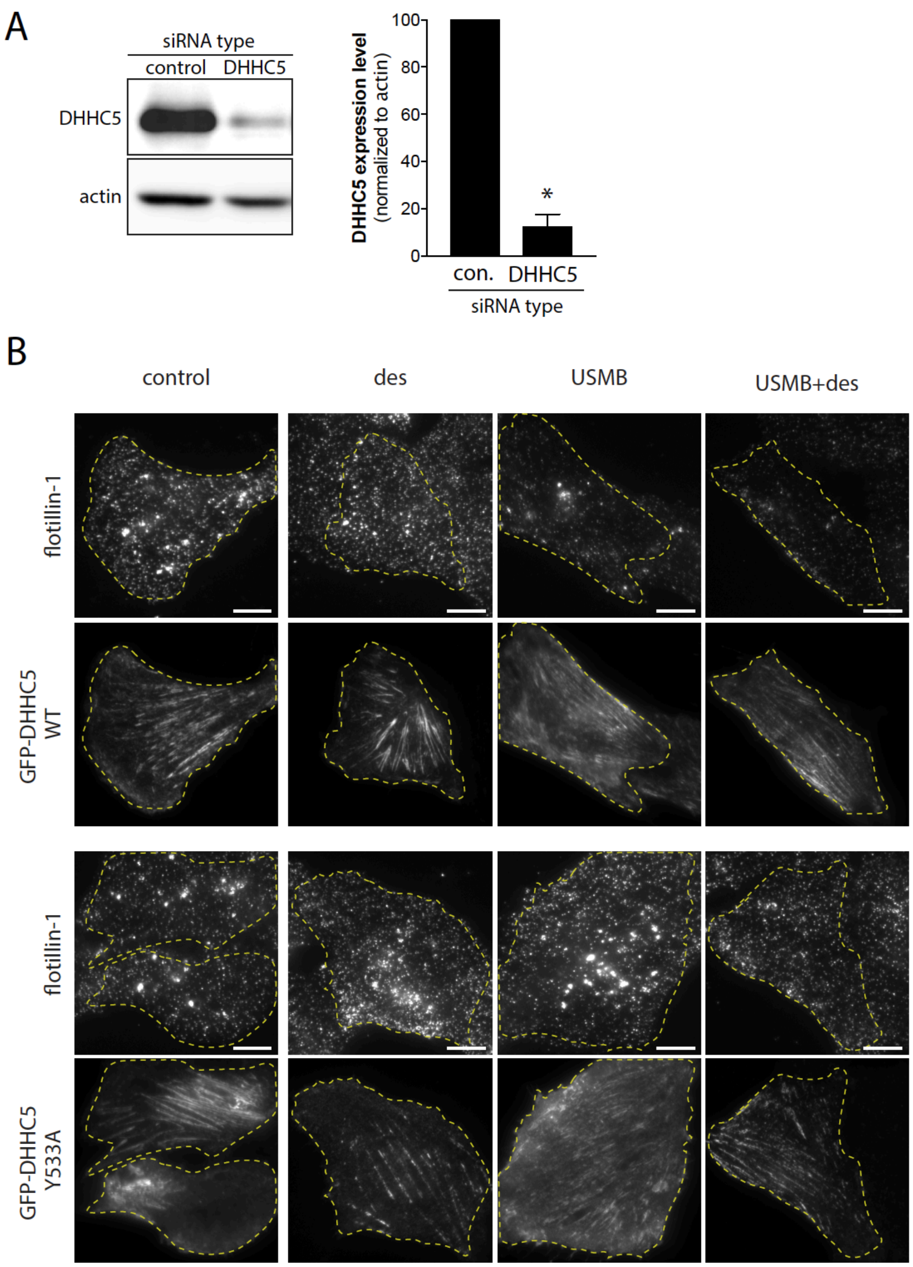


**Figure S2. DHHC5 knockdown and transfection**. (A) RPE cells were transfected with siRNA targeting DHHC5 or non-targeting siRNA (control). Whole-cell lysates were resolved by SDS-PAGE and subjected to immunoblotting to detect DHHC5 or actin (loading control). Shown (left panels) are representative immunoblot and the mean ± SE of DHHC5 detected by this method (right panels). This demonstrates effective knockdown of DHHC5 by siRNA silencing. (B) RPE cells were transfected with cDNA encoding either wild-type (WT) or Y533A mutant DHHC5, fused to eGFP. Following transfection, some cells were treated with 50 μM desipramine for 60 min followed by USMB treatment, as indicated. Following a subsequent 30 min incubation, cells were then fixed and subjected to immunofluorescence staining of flotillin-1, and then imaging by TIRF-M. Shown are representative TIRF-M fluorescence micrographs, scale 10 μm. Transfected cells are outlined, based on GFP signal. These images show the eGFP-DHHC5 expression of cells shown in **Figure 5**.

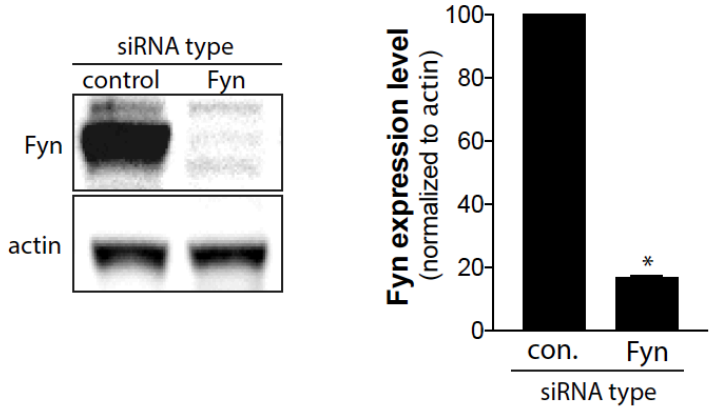


**Figure S3. Fyn knockdown.** RPE cells were transfected with siRNA targeting Fyn or non-targeting siRNA (control). Whole-cell lysates were resolved by SDS-PAGE and subjected by immunoblotting to detect Fyn or actin (loading control). Shown (left panels) are representative immunoblot and the mean ± SE of Fyn detected by this method (right panels). This demonstrates effective knockdown of Fyn by siRNA silencing.

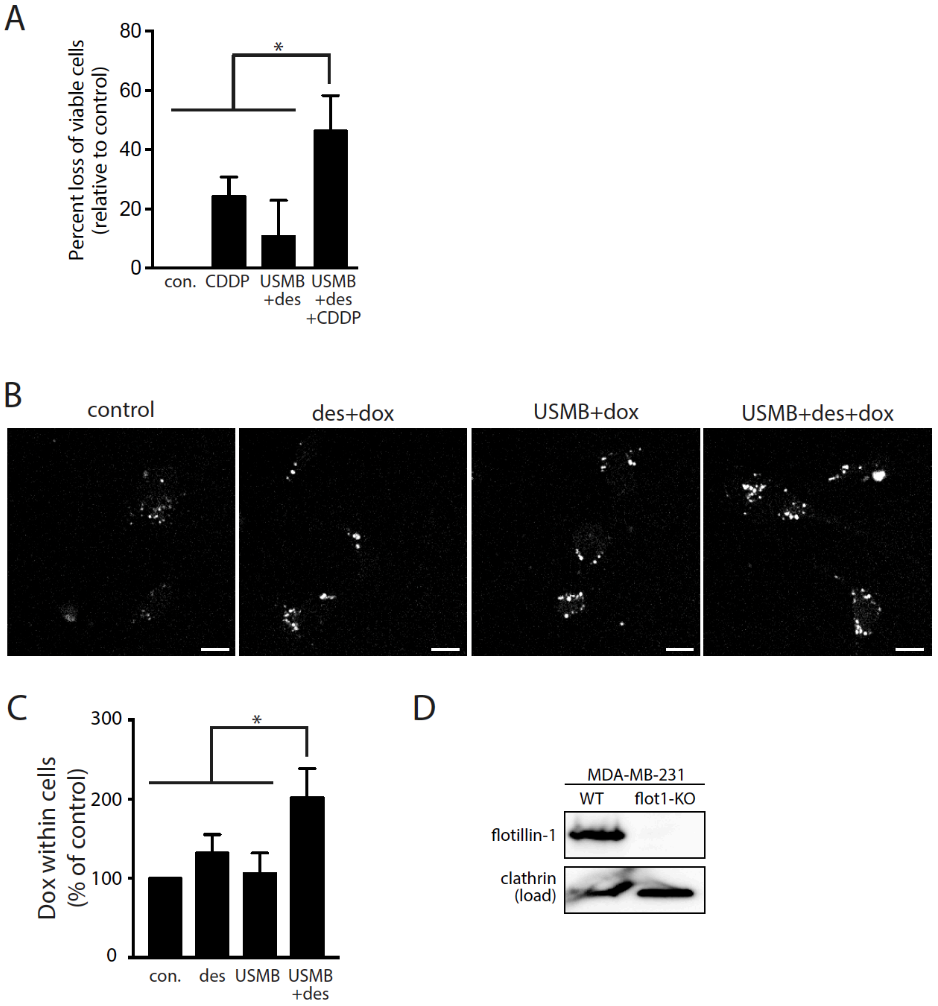


**Figure S4. Flotillin knockout cells and contribution of flotillin to cell viability in USMB-treated cells.** (A) RPE cells were treated with 50 $\mu$M desipramine for 60 min followed by USMB treatment, as indicated. Following treatments, some cells were incubated with 0.03 mM cisplatin (CDDP) for 2 h, as indicated, followed by washing and incubation in growth media (no drugs). 24 h after USMB and/or cisplatin exposure, cell viability was assessed by crystal violet assay, and shown are the mean ± SE of the percent reduction in the total viable cells, relative to that of control (not treated with CDDP, USMB or desipramine). n = 3 independent experiments. *, $p < 0.05$. (B-C) RPE cells were treated with 50 $\mu$M desipramine for 60 min followed by USMB treatment, as indicated. Following treatments, cells were incubated with 0.03 mM doxorubicin for 2 h (as shown), followed by washing and incubation in growth media (no drugs) for 24 h, after which cells were subjected to widefield epifluorescence microscopy to detect doxorubicin fluorescence within cells. Shown in (B) are representative fluorescence micrographs of doxorubicin fluorescence, and in (C) the mean ± SE of cellular doxorubicin fluorescence. n = 3 independent experiments. *, $p < 0.05$. This indicates that cells treated with USMB+desipramine retain more chemotherapeutic drugs 24 h after exposure to these drugs compared to control cells (cells not treated with USMB+desipramine). (D) Whole cell lysates from MDA-MB-231 wild-type (WT) cells or MDA-MB-231 cells subjected to CRISR/Cas9 genome editing (to knock out flotillin-1) were resolved by SDS-PAGE and subjected by immunoblotting to detect flotillin-1 or clathrin heavy chain (loading control). Shown are representative immunoblots demonstrates effective knockout of flotillin in MDA-MB-231-flot-KO cells.
